# Altered hippocampal transcriptome dynamics following sleep deprivation

**DOI:** 10.1101/2021.05.20.445021

**Authors:** Marie E. Gaine, Ethan Bahl, Snehajyoti Chatterjee, Jacob. J. Michaelson, Ted Abel, Lisa C. Lyons

**Author notes:** **Corresponding Author**: Lisa C. Lyons.

## Abstract

Widespread sleep deprivation is a continuing public health problem in the United States and worldwide affecting adolescents and adults. Acute sleep deprivation results in decrements in spatial memory and cognitive impairments. The hippocampus is vulnerable to acute sleep deprivation with changes in gene expression, cell signaling, and protein synthesis. Sleep deprivation also has long lasting effects on memory and performance that persist after recovery sleep, as seen in behavioral studies from invertebrates to humans. Although previous research has shown that acute sleep deprivation impacts gene expression, the extent to which sleep deprivation affects gene regulation remains unknown. Using an unbiased deep RNA sequencing approach, we investigated the effects of acute sleep deprivation on gene expression in the hippocampus. We identified 1,146 genes that were significantly dysregulated following sleep deprivation with 507 genes upregulated and 639 genes downregulated, including protein coding genes and long non-coding RNAs not previously identified as impacted by sleep deprivation. Notably, genes significantly upregulated after sleep deprivation were associated with RNA splicing and the nucleus. In contrast, downregulated genes were associated with cell adhesion, dendritic localization, the synapse, and postsynaptic membrane. These results clearly demonstrate that sleep deprivation differentially regulates gene expression on multiple transcriptomic levels to impact hippocampal function.

**Graphical Abstract:** 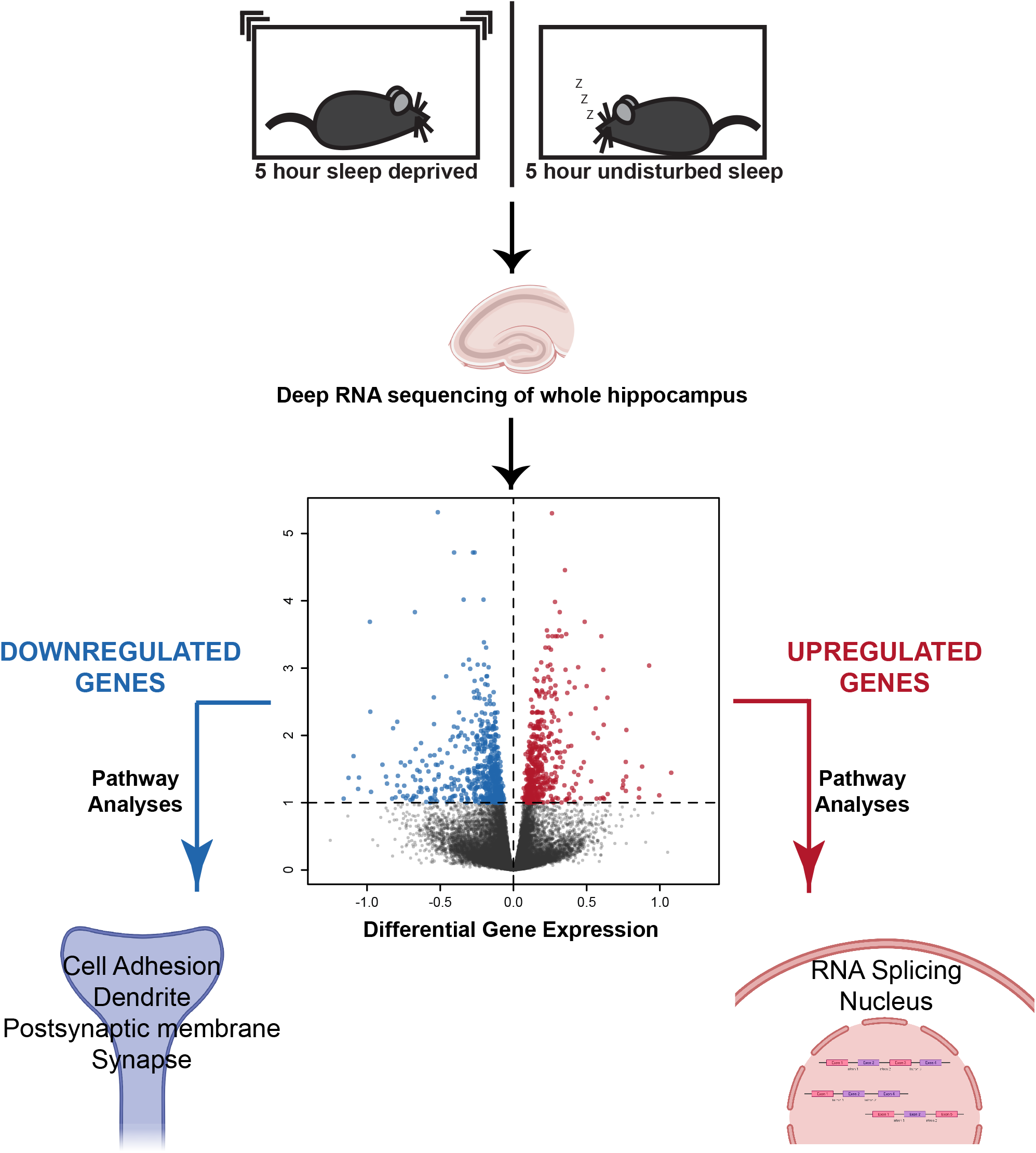

## Introduction

Sleep deprivation is a widespread public health problem in the United States and countries around the globe [1]. In the United States, estimates suggest that nearly 70% of adults and teenagers have insufficient sleep at least one day per month [2-4]. Acute sleep deprivation results in cognitive impairments (reviewed in [5]), as well as the exacerbation of neuropsychiatric and mood disorders (reviewed in [6, 7]). The decrements in cognitive function and performance induced by acute sleep deprivation create an economic burden with decreased workplace productivity as well as increased accident risk encumbering public safety [8-11]. Moreover, acute sleep deprivation results in increased levels of amyloid-beta as well as increased levels of tau in cerebral spinal fluid and plasma, which are pathological markers associated with increased risk of Alzheimer’s disease [12, 13].

The impact of sleep deprivation on long-term memory is phylogenetically conserved as seen in behavioral studies of invertebrates to rodent models to human subjects [14-18]. Moreover, the effects of acute sleep deprivation on memory can extend for days, even with recovery sleep. For example, in the marine mollusk *Aplysia*, the effects of acute sleep deprivation persist for at least 48 hours inhibiting the formation of long-term memory [17]. Similarly, in humans, acute deprivation impairs episodic memory and hippocampus dependent memory associations for more than two days, despite recovery sleep [18]. Long-lasting cellular mechanisms such as changes in gene regulation ostensibly underlie the conserved persistent effects of acute sleep deprivation on memory. The hippocampus is particularly susceptible to the impacts of sleep deprivation with changes apparent in cellular signaling, protein synthesis, and neuronal connectivity following sleep deprivation [12, 18-22], although studies differ as to the effect of acute sleep deprivation on dendritic structure [13, 22-24]. Studies have highlighted the effects of sleep deprivation on gene expression and protein synthesis in the hippocampus [25-27], with more recent analyses conducted to identify gene networks and hubs correlated with sleep deprivation [28]. Sleep deprivation also induces epigenetic alterations affecting gene expression in animal models and humans ([29, 30] and reviewed in [31]). Enhancement of global gene transcription through inhibition of histone deacetylation has been shown to rescue hippocampus-dependent memory and synaptic plasticity in sleep deprived mice [32]. Thus, our understanding of the extent and specificity of sleep deprivation on gene regulation remains incomplete.

To more fully detail the effects of acute sleep deprivation on transcription, we investigated the effect of 5 hours of sleep deprivation on gene expression in the hippocampus using an unbiased deep RNA sequencing (RNA-Seq) approach. We identified 1,146 genes differentially regulated after sleep deprivation. Genes significantly upregulated were preferentially associated with the nucleus with functions in RNA binding and processing, whereas genes significantly downregulated after sleep deprivation were associated with cell adhesion, the synapse, dendrites, and postsynaptic membrane. Through comparison with a recently published data set analyzing the effects of acute sleep deprivation on ribosome associated transcripts in excitatory neurons of the hippocampus [27], we found a considerable difference between the number of genes regulated by sleep deprivation at the total RNA level in the ribosome associated pool of transcripts. Genes regulated by sleep deprivation at both the transcriptional and translational levels showed enrichment in protein kinase and phosphatase activity, as well as potassium and cation channel activity. Functions enriched with genes regulated by sleep deprivation only in the transcriptome included transcription factor binding, histone deacetylase activity, nucleotide binding, nucleotide exchange factor activity and small GTPase regulator activity; whereas genes regulated solely in the translatome displayed network enrichment for the unfolded protein binding pathway, protein binding, peptide binding, protein dimerization and ubiquitin binding. The data set generated with this research highlights the differences in biological function between genes upregulated after sleep deprivation and those downregulated demonstrating the gene specific effects of sleep deprivation and recovery sleep on gene regulation.

## Materials and Methods

### Animals

C57BL/6J (Jackson Labs, #000664) male mice between 3-4 months of age were used for all experiments. Mice were housed in groups of up to five under 12 hour light/12 hour dark cycle with *ad libitum* access to food and water in a temperature and humidity controlled room (22°C and 55 +/**-** 5%, respectively). Mice were maintained under standard conditions consistent with National Institute of Health guidelines and approved by the Institutional Animal Care and Use Committee of the University of Iowa.

### Sleep deprivation and recovery

One week prior to sleep deprivation mice were single housed with corn cob bedding as previously described [25-27]. Three days prior to sleep deprivation, mice were gently handled for 3 minutes, with cages lightly tapped or moved. Sleep deprivation was performed starting at the beginning of the light cycle to control for circadian differences in gene expression. Sleep deprivation was carried out for 5 hours using the gentle handling method [14, 25]. Animals were monitored continuously with minimal cage tapping and then cage shakes as necessary disturbances to achieve sleep deprivation. Non-sleep deprived mice were placed in a room by themselves with lighting and humidity similar to the sleep deprivation room and left undisturbed for 5 hours. To study sleep recovery, mice were sleep deprived for 5 hours and then returned to the housing room for 3 hours to recover. Control animals were left undisturbed. For all experiments, control and experimental animals were sacrificed at the same time to avoid circadian confounds. At the end of undisturbed sleep, sleep deprivation, or recovery, hippocampi were removed and flash frozen.

### RNA extraction

Hippocampal samples were homogenized in Qiazol (Invitrogen) and phase separated using chloroform followed by centrifugation at 14,000 g for 15 minutes. RNA was extracted using the RNeasy kit (Qiagen) with DNA removed with RNase-Free DNase (Qiagen). Samples were resuspended in RNase-free water and quantified using the Nanodrop 1 and the Agilent Bioanalyzer. Samples with an OD 260/280 and OD 260/230 ratio close to 2.0 and RNA integrity number (RIN) above 8 were selected for library preparation.

### RNA library preparation and sequencing

RNA libraries were prepared at the Iowa Institute of Human Genetics, Genomics Division using the Illumina TruSeq Stranded Total RNA with Ribo-Zero gold sample preparation kit (Illumina). Library concentrations were measured with the KAPA Illumina Library Quantification Kit (KAPA Biosystems). Pooled libraries were sequenced across two lanes in 150 bp paired-end reads using the Illumina HiSeq 4000. A total of 18 samples were sequenced in two batches.

### RNA Sequencing analysis

The bcbio-nextgen pipeline [33] was used to process sequencing data. STAR [34] was used to align reads to the mm10 genome build and featureCounts was used to quantify expression at the gene level [35]. EDASeq was used to adjust for GC content effects and account for sequencing depth [36] with normalization shown in Additional Figure S1. Run length encoding (RLE plots) and PCA analysis were used to validate normalization (Additional Figure S1). Differential expression analysis was conducted using edgeR’s quasi-likelihood pipeline [37-39]. Effect size was calculated by removing gene abundance from fold changes. The RNA-Seq data have been deposited in NCBI’s Gene Expression Omnibus and are accessible through GEO Series accession number GSE166831, https://www.ncbi.nlm.nih.gov/geo/query/acc.cgi?acc=GSE166831. The code for analyses and figures related to RNA-Seq data can be accessed through GitHub at https://github.com/ethanbahl/gaine2021_sleepdeprivation.

### Pathway Analysis

To identify biological pathways affected by sleep deprivation, we used NetworkAnalyst V3.0 [40] to perform Global Enrichment Network OverRepresentation Analyses (ORA) on upregulated and downregulated genes (< 0.1 FDR) using the Protein ANalysis THrough Evolutionary Relationships (PANTHER):BP and PANTHER:CC classifications to determine biological processes (BP) and cellular components (CC) [41]. For the pathway analyses, a P-value < 0.05 was considered significant. Comparison of the RNA-seq data from this manuscript with the TRAP-Seq data available in NCBIs GEO repository, GEO series accession GSE156925, https://www.ncbi.nlm.nih.gov/geo/query/acc.cgi?acc=GSE156925, [27], was performed using Gene Ontology-Molecular Function (GO:MF) due to the relatively small number of differentially expressed genes in common between the two data sets.

### Quantitative real-time PCR (RT-qPCR)

RNA (500 ng) was used for cDNA preparation with the Superscript IV First-Strand Synthesis (Ambion). RT-qPCR was performed with gene specific primers (Additional File 2: Table S1) using Fast SYBR™ Green Master Mix (ThermoFisher Scientific). Reactions were run using the Quant Studio 7 Real-Time PCR System (ThermoFisher Scientific). Samples were quantified in at least triplicate using two appropriate housekeeping genes (*Tubulin* and *Hprt*) that were included on all plates for normalization. Neither housekeeping gene was differentially expressed between non-sleep deprived and sleep deprived samples in the RNA-seq data set: *Tuba1a* Log Fold Change (FC) = 0.018, FDR = 0.882; and *Hprt* LogFC = −0.003, FDR = 0.981.

### Statistical Analyses

RT-qPCR statistical analyses were performed using an unpaired Student’s t-test in GraphPad Prism. Results were expressed as means +/- SEM. Values of P < 0.05 were considered as statistically significant.

## Results

### Deep RNA sequencing reveals the extent of changes in gene expression induced in the hippocampus by acute sleep deprivation

Previously researchers analyzed the impact of sleep deprivation on gene expression in the hippocampus using microarrays [25]; however, this approach had limitations in detection due to the microarray chip design, i.e., probes must be designed *a priori* that target specific anticipated transcripts. In contrast, RNA-seq provides an unbiased approach to identify differential gene expression rather than relying on a set of predetermined sequences. Moreover, deep RNA sequencing facilitates identification of differential expression for non-coding sequences including antisense transcripts, long non-coding RNAs and microRNAs [42, 43]. Accordingly, we investigated the effects of 5 hour acute sleep deprivation starting at lights on using gentle handling to identify changes in gene expression in the hippocampus with deep RNA sequencing. Sleep deprivation experiments (Figure 1a) were completed in two independent cohorts (cohort one had n=4 and cohort two had n=5 samples for each of the non-sleep deprived and sleep deprived group). Through deep sequencing with an average of 143M reads per sample in the non-sleep deprived mice and 136.5M reads per sample in the sleep deprived mice, we found sufficient expression levels for 22,582 genes including protein coding, non-coding, and predicted genes. GC and depth normalization were used to normalize the raw data before processing (Additional File 1: Figure S1).

**Figure 1.**
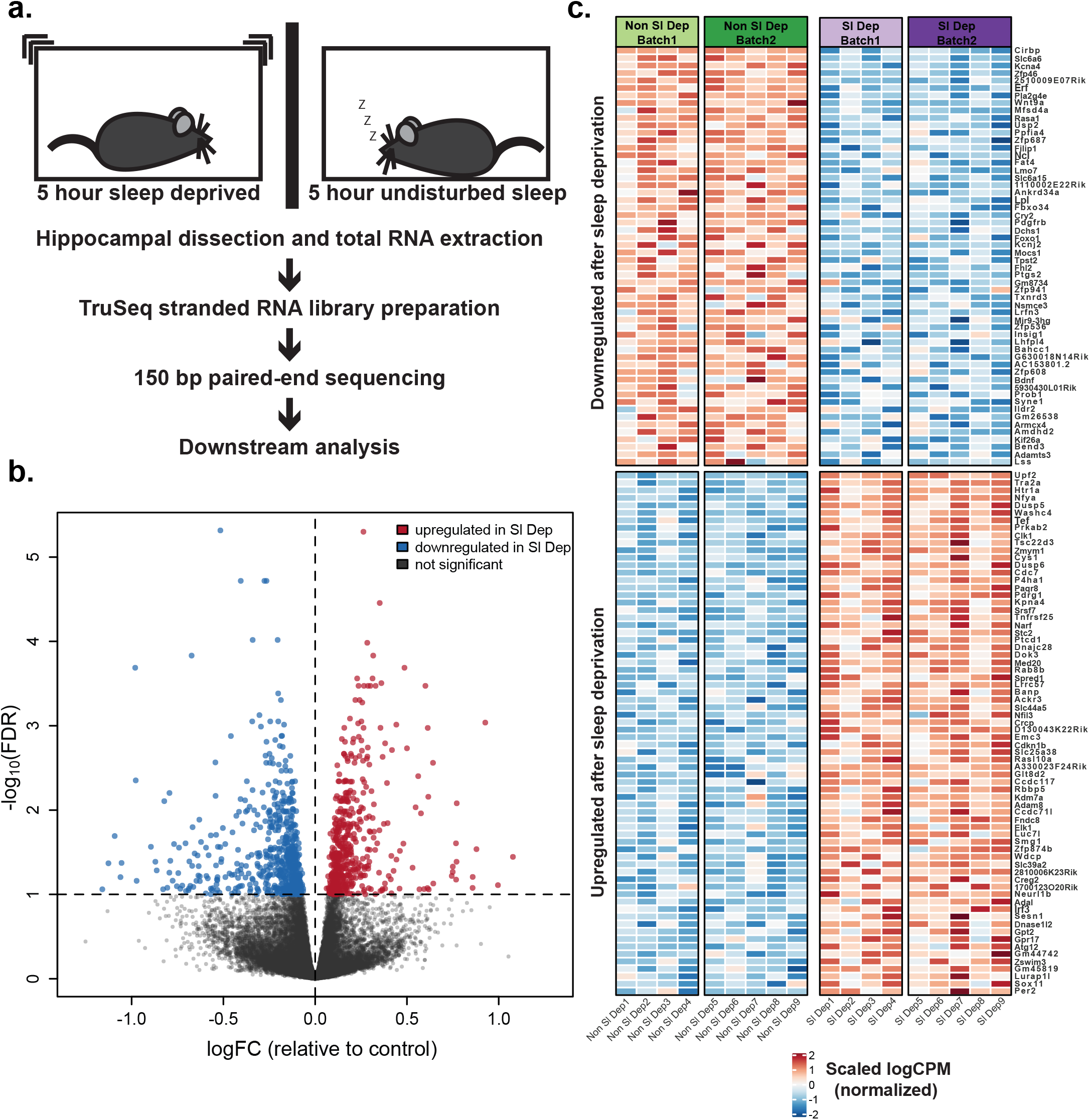
Acute sleep deprivation causes substantial gene expression changes in the mouse hippocampus. **(a)** Schematic showing experimental procedure for RNA sequencing following acute sleep deprivation. C57BL/6J male mice were either sleep deprived for 5 hours (n=9) or left undisturbed (n=9). Immediately following sleep deprivation or undisturbed sleep, the whole hippocampus was dissected out and flash frozen. Total RNA was extracted and processed for RNA sequencing. **(b)** Volcano plot illustrating differentially expressed genes between non-sleep deprived and sleep deprived mice. Genes with a false discovery rate (FDR) < 0.1 are highlighted in red for significantly upregulated (507 genes) and blue for significantly downregulated (639 genes) after sleep deprivation. Genes that are not significantly differentially expressed in sleep deprived mice are in grey. **(c)** Heatmap showing the most differentially expressed genes filtered by FDR ≤ 0.01 and effect size > ±0.5 in each cohort. The top rows represent genes that are significantly downregulated after sleep deprivation. The bottom rows represent genes that are significantly upregulated after sleep deprivation. Each column represents one mouse and columns are grouped by batch. The scale represents log counts per million (logCPM), with red denoting upregulation and blue denoting downregulation after sleep deprivation. The most significantly upregulated gene after sleep deprivation was UPF2 Regulator of Nonsense Mediated MRNA Decay (*Upf2*; log fold change (LogFC) = 0.263, FDR = 5.01 × 10^− 6^). The most significantly downregulated gene after sleep deprivation was Cold Inducible RNA Binding Protein (*Cirbp*; LogFC = −0.516, FDR = 4.83 × 10^−6^).

We identified 1,146 genes that were significantly dysregulated following sleep deprivation (FDR < 0.10) with 507 genes upregulated and 639 genes downregulated (Figure 1b and Additional File 2: Tables S2 and S5). Of the differentially expressed genes, 1,026 (89.5%) were protein coding based on the Ensembl biotype classification [44]. Heatmap representations are shown for differentially regulated genes based on FDR (≤ 0.01) and effect size (absolute value > ±0.5) to identify those genes most strongly affected by sleep deprivation (Figure 1c). The most significantly upregulated gene after sleep deprivation was a Regulator of Nonsense Mediated mRNA Decay (*Upf2*), while the most significantly downregulated gene after sleep deprivation was Cold Inducible RNA Binding Protein (*Cirbp*). In these experiments, the animals were sleep deprived starting at lights on, while in the microarray study by Vescey and colleagues, sleep deprivation was initiated four to six hours after lights on; however, *Upf2* and *Cirbp* were differentially expressed after sleep deprivation in both studies. Using a Fisher’s Exact Test to compare the overlap between the RNA-Seq differential gene expression and the microarray gene expression, we found strong overlap between the data sets with an odds ratio of 8.43 (confidence interval [7.05, 10.06]; P-value < 2.2 × 10^−16^). We identified 226 differentially regulated genes in common between the two data sets, suggesting that sleep deprivation targets a core set of genes regardless of whether sleep deprivation begins at the beginning of the rest period or after a few hours of rest.

With the *de novo* sequencing approach, we found 6,967 sequences that were not previously tested in the microarray study by Vecsey and colleagues [25]. Using Ensembl biotype classification, 973 genes that were previously not analyzed with sleep deprivation were determined to be protein coding. Within this additional gene set, there were 123 sequences that were significantly differentially expressed between non-sleep deprived and sleep deprived groups (FDR < 0.10) including 33 protein coding genes, 16 long intergenic non-coding RNAs (lincRNAs), 2 long non-coding RNAs, 1 microRNA, 14 pseudogenes, 13 antisense transcripts, 39 EST sequences, and 5 processed transcripts. Thus, unbiased deep RNA sequencing provided a more thorough analysis of differential gene expression than previous research using microarrays.

### RNA splicing and nuclear localization are associated with genes upregulated after sleep deprivation

To identify biological and functional relevance, we separately analyzed the upregulated and downregulated genes after acute sleep deprivation using Network Analyst and the PANTHER:BP classification to perform network enrichment. For genes significantly upregulated by sleep deprivation, there were 16 pathways significantly enriched (Figure 2a; Additional File 2: Table S3). Notably, RNA splicing was one of the most significant pathways identified with genes upregulated after sleep deprivation. As shown in Figure 2a, the enrichment of RNA splicing and processing pathways included 14 genes upregulated after sleep deprivation. We also found enrichment for pathways involved in circadian rhythms and rhythmic processes highlighting the interactions between sleep deprivation and the circadian clock. To predict the cellular localization of proteins for which sleep deprivation induced increased gene expression, we utilized the PANTHER:CC classification (Figure 2b and Additional File 2: Table S4). Genes upregulated after sleep deprivation were most commonly associated with nuclear localization, consistent with the hypothesis that processes critical to RNA splicing and RNA processing are impacted by acute sleep deprivation.

**Figure 2.**
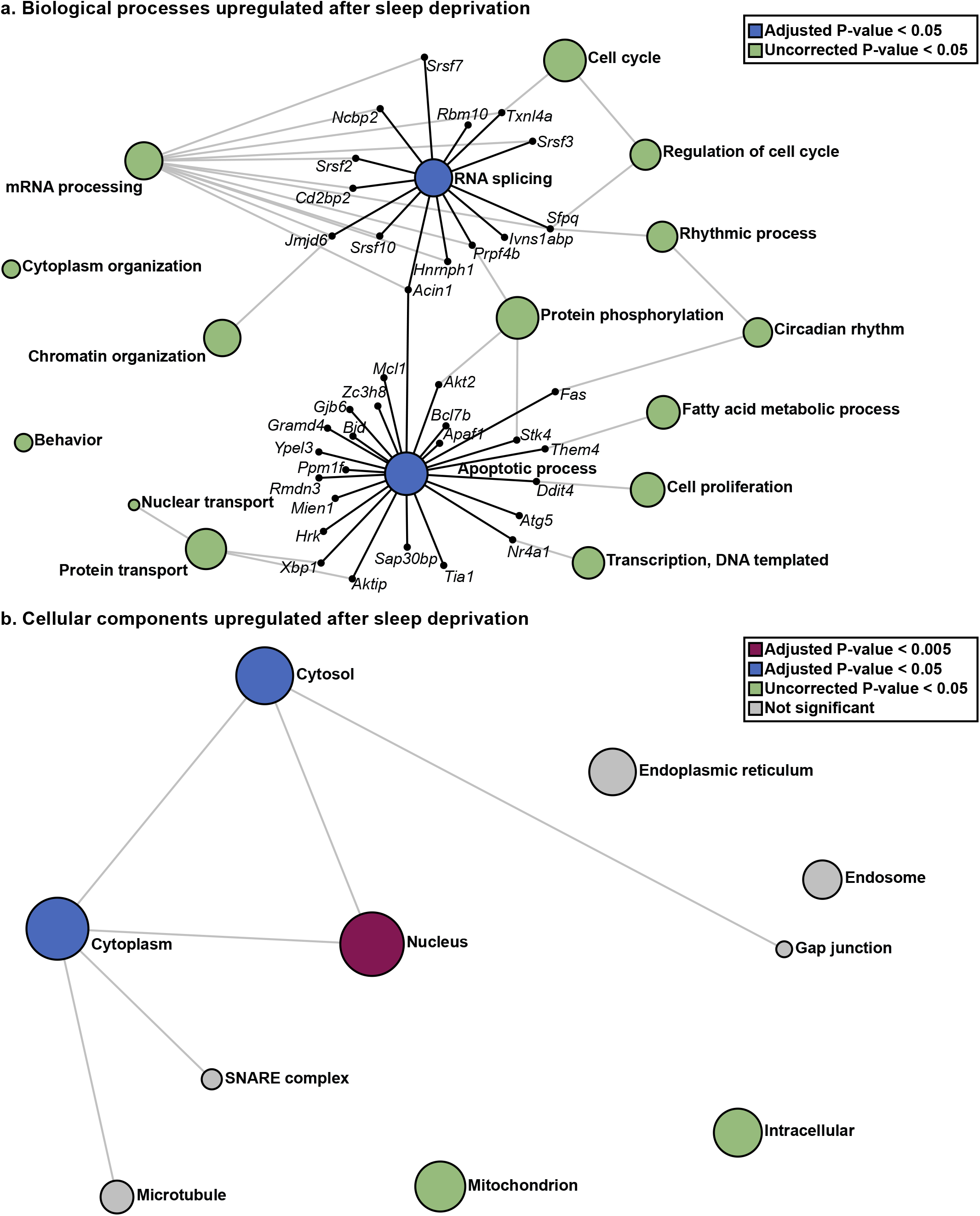
Distinct biological processes and cellular components are enriched for genes upregulated after acute sleep deprivation. **(a)** Using Network Analyst software and the PANTHER: Biological Processes (BP) classification to perform overrepresentation analysis (ORA), pathways enriched for upregulated genes were identified. RNA splicing and Apoptotic Process were the most significantly enriched networks (Adjusted P-value = 0.025). These top networks have been expanded to show the genes that are involved and upregulated after sleep deprivation. **(b)** Using Network Analyst software and the PANTHER: Cellular Components (CC) classification to perform ORA we identified cellular components enriched for genes upregulated after sleep deprivation. The nucleus is the most significantly enriched (Adjusted P-value = 1.74 × 10^−9^). The size of each node represents the number of hits from the inputted gene list.

### Genes downregulated by sleep deprivation are associated with dendritic, postsynaptic membrane, and cytoskeletal components

Analysis of the genes significantly downregulated after sleep deprivation revealed strikingly different biological functions and cellular localization than those genes that were upregulated after sleep deprivation. When analyzing network enrichment using PANTHER:BP, we identified 19 pathways enriched for genes downregulated after sleep deprivation. The most significantly enriched pathway was Cell Adhesion (Figure 3a and Additional File 2: Table S6), with this pathway including 31 genes significantly downregulated after sleep deprivation. In contrast to the strong nuclear association seen for genes upregulated by sleep deprivation, analysis using PANTHER:CC revealed that the genes downregulated by sleep deprivation are associated with many different cellular components (Figure 3b and Additional File 2: Table S7). Dendrites, postsynaptic membranes, and the synapse were the three cellular compartments most significantly associated with genes downregulated after sleep deprivation. Downregulated genes were also significantly associated with the cytoplasm and cytoskeletal components. These results strongly suggest that sleep deprivation differentially regulates genes, at least in part, based on their biological function, rather than a more global regulation of gene expression.

**Figure 3.**
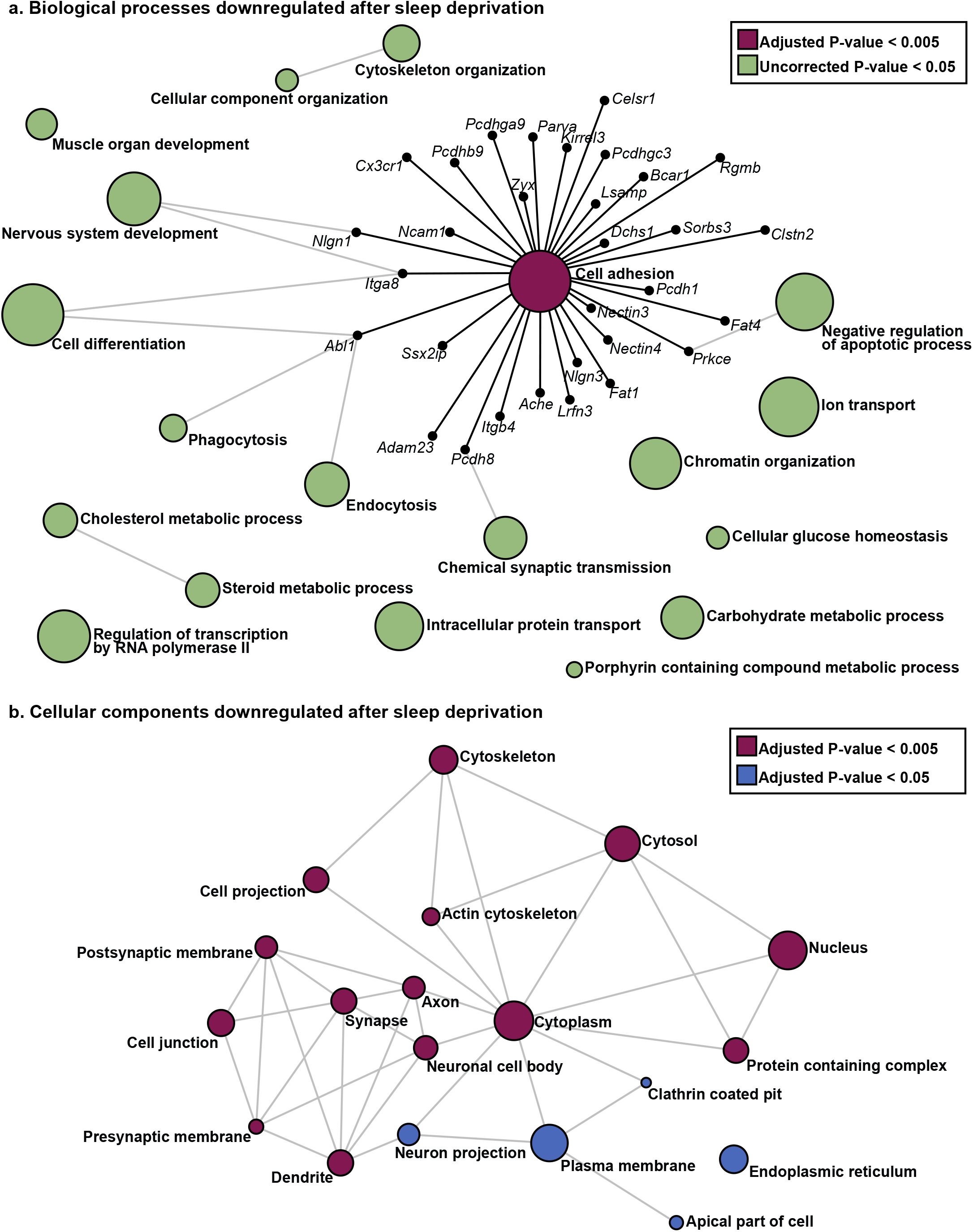
Distinct biological processes and cellular components are enriched for downregulated genes after acute sleep deprivation. **(a)** Pathway analysis using the PANTHER:BP classification to perform ORA, pathways enriched for downregulated genes were identified. Cell adhesion is the most significantly enriched network (Adjusted P-value = 2.91 × 10^− 3^). The cell adhesion network has been expanded to show the genes that are involved and downregulated after sleep deprivation. **(b)** Using Network Analyst software and the PANTHER:CC classification to perform ORA we identified enriched cellular components for genes downregulated after sleep deprivation. The dendrite and postsynaptic membrane are the most significantly enriched cellular components (Adjusted P-value = 4.28 × 10^−7^). The size of each node represents the number of hits from the inputted gene list.

### Confirmation of gene expression changes in the hippocampus after sleep deprivation

Based upon molecular function, we chose a subset of the differentially expressed genes from the RNA-Seq experiment to validate in independent sleep deprivation experiments (Figure 4; non-sleep deprived mice n=6, sleep deprived mice n=6). Given the significance of pathways involved in RNA processing including RNA splicing, we tested the effects of sleep deprivation on the expression of four genes, *Cirbp, Srsf7, Tra2a*, and *Upf2*, with known functions in RNA processing or as RNA binding proteins (Figure 4a). Using RT-qPCR with gene specific primers (Additional File 2: Table S1), we confirmed that these genes demonstrated significant changes with the same directionality after acute sleep deprivation. Immediate early genes and transcription factors have been previously identified as changing after acute sleep deprivation in the hippocampus and other brain regions [45-50]. We independently confirmed that acute sleep deprivation differentially regulated two positive transcription factors and a repressor of transcription, *Nfil3, Nr4a1* and *Erf* (Figure 4b). We also validated through RT-qPCR four genes involved in cellular signaling that were significantly changed after sleep deprivation, *Pdgfrb, Dusp5, Dusp6*, and *Ackr3* (Figure 4c). Acute sleep deprivation has been previously shown to induce changes in the cytoskeleton and decrease dendritic spines in the CA1 and dentate gyrus of the hippocampus [20, 22]. We validated the effects of sleep deprivation on two genes of interest that act in the postsynaptic dendrites, *Filip1* and *Arc* (Figure 4d). All the genes included were significantly changed in the independent cohort.

**Figure 4.**
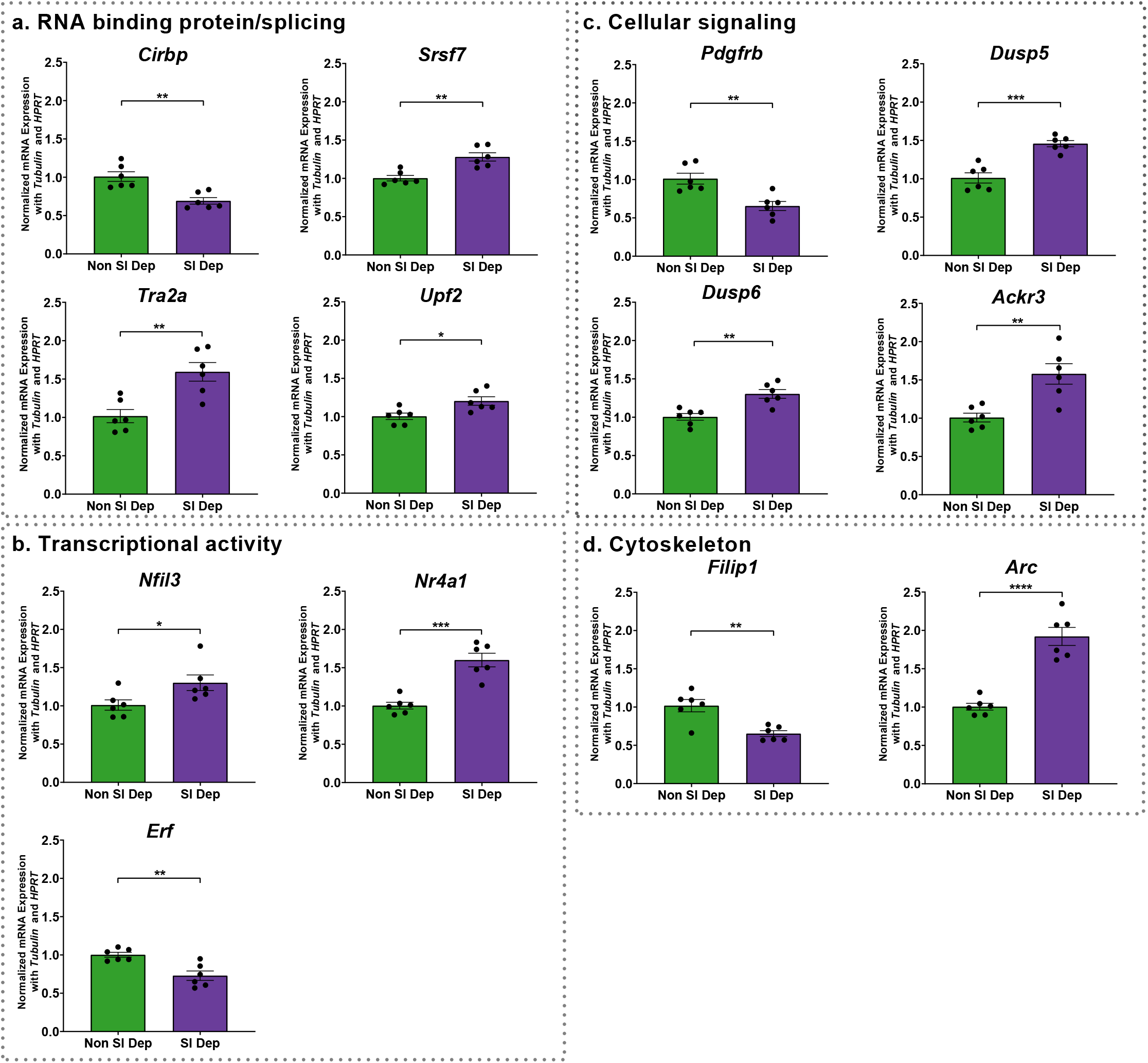
RT-qPCR validation of RNA sequencing results. From an independent cohort of mice (n=6 in each group), RT-qPCR was used to validate the findings of chosen genes. **(a)** Four genes related to RNA binding proteins and/or splicing: *Cirbp* (P-value = 1.9 × 10^−3^), *Srsf7* (P-value = 1.6 × 10^−3^), *Tra2a* (P-value = 3.0 × 10^−3^), and *Upf2* (P-value = 0.0168); **(b)** three genes related to transcriptional activity: *Nfil3* (P-value = 0.0388), *Nr4a1* (P-value = 1.0 × 10^−4^), and *Erf* (P-value = 2.6 × 10^−3^); **(c)** four genes related to cellular signaling: *Pdgfrb* (P-value = 3.2 × 10^−3^), *Dusp5* (P-value = 2.0 × 10^−4^), *Dusp6* (P-value = 2.0 × 10^−3^), and *Ackr3* (P-value = 2.8 × 10^−3^); and **(d)** two genes related to cytoskeleton: *Filip1* (P-value = 2.1 × 10^−3^) and *Arc* (P-value < 0.0001). Data are presented as mean ± SEM and normalized against two housekeeping genes (*Tubulin* and *Hprt*). Differences are significant at * P ≤ 0.05, ** P ≤ 0.01, *** P ≤ 0.001, **** P < 0.0001, and evaluated using an unpaired t-test.

As a negative control to validate our analysis, we chose three genes with varying functions that showed no significant difference in expression after sleep deprivation in the previous microarray study (GEO accession GSE33302) or between non-sleep deprived and sleep deprived samples in our RNA-Seq data set: Laminin Subunit Alpha 5 (*Lama5*), Frizzled Class Receptor 5 (*Fzd5*), and Transient Receptor Potential Cation Channel Subfamily M Member 3 (*Trpm3*). Using RT-qPCR, we quantified the expression of these three genes after sleep deprivation from the independent cohort of animals. As predicted, we did not find significant differences in the expression levels of these genes between non-sleep deprived and sleep deprived animals (Additional File 1: Figure S2).

### Three hours of sleep recovery reverses the effects of acute sleep deprivation on mRNA abundance

Previous studies have shown that 2.5 – 3 hours of recovery sleep following acute sleep deprivation is sufficient to reverse many of the effects of sleep deprivation on gene expression, cellular signaling that affects protein synthesis, and dendritic structure [22, 25, 26, 51, 52]. However, recovery sleep following acute sleep deprivation is unable to restore the deficits observed in long-term hippocampus dependent memory [15]. Consequently, we investigated whether recovery sleep following acute sleep deprivation was sufficient to normalize mRNA abundance for a subset of genes, including those associated with synaptic plasticity and memory (Figure 5). We performed sleep deprivation experiments followed by 3 hours of recovery sleep. Control animals were euthanized at the same time as experimental animals to avoid circadian confounds in gene expression (n=6 for non-sleep deprived mice and n=7 for sleep deprived with recovery sleep group). For most genes, we found that 3 hours of recovery sleep induced a return to baseline gene expression levels with no significant changes when sleep deprived plus recovery sleep mice were compared with non-sleep deprived mice: *Cirbp, Tra2a, Upf2, Nfil3, Erf, Pdgfrb, Dusp5, Dusp6, Ackr3*, and *Filip1*. Notably, the transcription factor *Nr4a1* remained significantly upregulated in recovery mice compared to non-sleep deprived mice; however, the fold change was significantly lower in the sleep recovery mice (mean fold change = 1.15) compared to that seen for gene expression after sleep deprivation (mean fold change = 1.60). Furthermore, recovery sleep significantly repressed mRNA abundance for the splicing factor *Srsf7*, a component of the spliceosome, that was upregulated after sleep deprivation. These results suggest that recovery sleep reverses acute sleep deprivation induced changes in gene expression for most genes, but exceptions occur for some genes which may affect RNA splicing or transcription, potentially providing a link to the more persistent effects of acute sleep deprivation.

**Figure 5.**
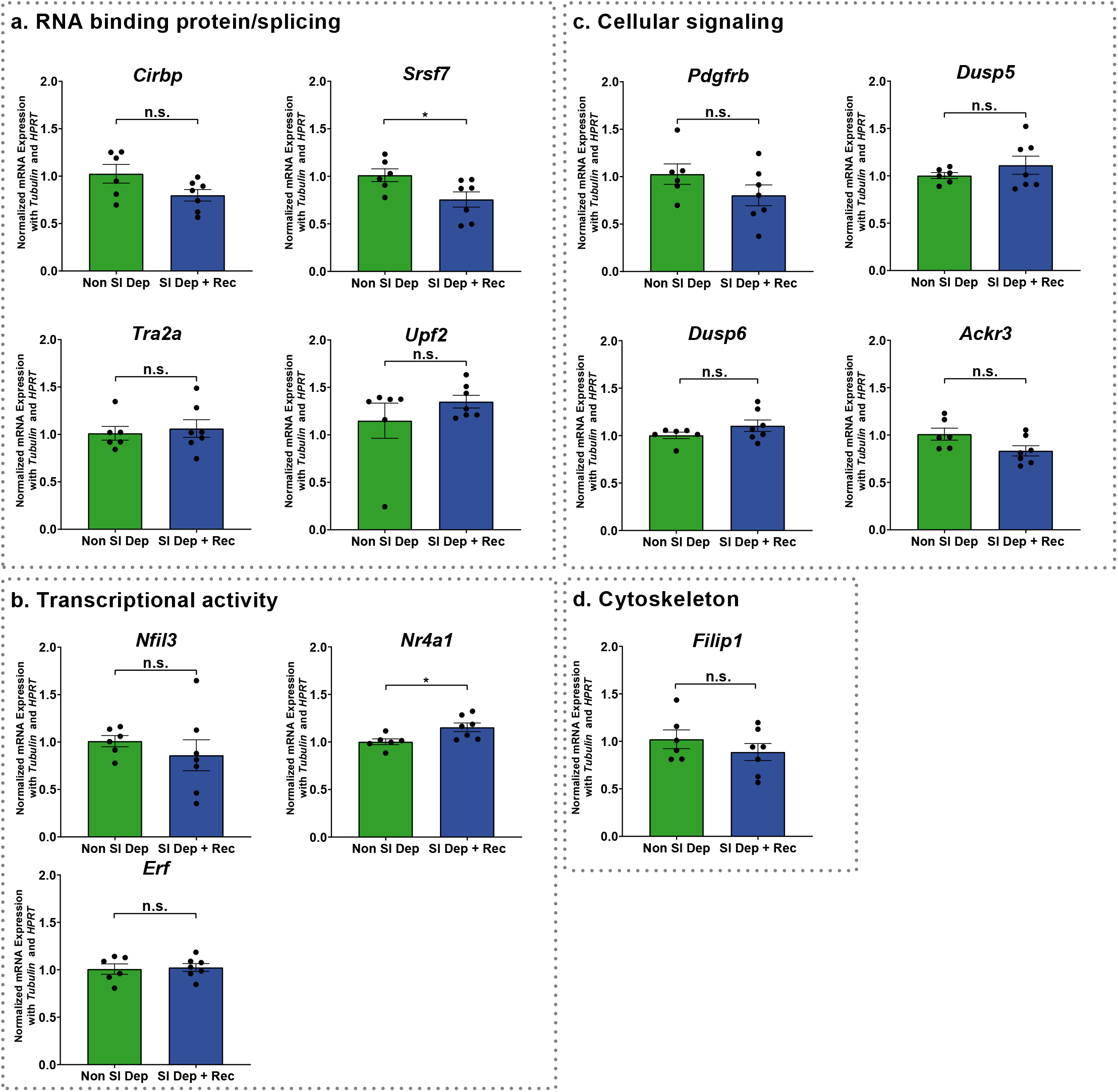
RT-qPCR analysis of chosen genes following recovery from sleep deprivation. An independent cohort of mice were allowed to recover from acute sleep deprivation for 3 hours (n=7) and compared to non-sleep deprived mice (n=6) using RT-qPCR analysis. **(a)** Four genes related to RNA binding proteins and/or splicing: *Cirbp* (P-value = 0.0674), *Srsf7* (P-value = 0.0356), *Tra2a* (P-value = 0.6881), and *Upf2* (P-value = 0.3009); **(b)** three genes related to transcriptional activity: *Nfil3* (P-value = 0.4402), *Nr4a1* (P-value = 0.0216), and *Erf* (P-value = 0.8060); **(c)** four genes related to cellular signaling: *Pdgfrb* (P-value = 0.1771), *Dusp5* (P-value = 0.3339), *Dusp6* (P-value = 0.1915), and *Ackr3* (P-value = 0.0565); and **(d)** one related to cytoskeleton: *Filip1* (P-value = 0.3342). Data are presented as mean ± SEM and normalized against two housekeeping genes (*Tubulin* and *Hprt*). Differences are significant at * P < 0.05 and evaluated using an unpaired t-test. n.s. denotes non-significant differences (P > 0.05).

### Acute sleep deprivation affects different gene functions in the transcriptome and translatome

Sleep deprivation can affect gene regulation at multiple levels with decreases in protein synthesis apparent in the hippocampus in addition to changes in mRNA abundance. Recent research investigated the effects of acute sleep deprivation on the pool of mRNA transcripts associated with ribosomes (TRAP-Seq) in excitatory neurons, which denotes the sleep deprived translatome [27]. To discriminate the effects of sleep deprivation on the transcriptome and the translatome, we performed a comparative analysis of the results from the current study with the translatome of excitatory neurons from published data (GEO Series accession GSE156925) using an FDR of < 0.10. Although both research studies used the same method of sleep deprivation performed at the same circadian time, we found that only 111 genes were similarly differentially regulated after sleep deprivation in both the transcriptome and translatome (Figure 6a and Additional File 2: Table S8). The limited overlap between the translatome and transcriptome is consistent with the limited overlap found when the translatome was previously compared to the microarray data set of Vecsey and colleagues [27]. As the TRAP-Seq data set is enriched for excitatory neurons, we predicted that the number of differentially expressed genes after sleep deprivation from the RNA-Seq data set would be greater as the RNA-Seq samples were prepared from the whole hippocampus including inhibitory neurons and glia. We found that 1,035 genes were differentially regulated in the transcriptome, but not in the translatome and 154 genes were regulated in the translatome but not in the whole transcriptome.

**Figure 6.**
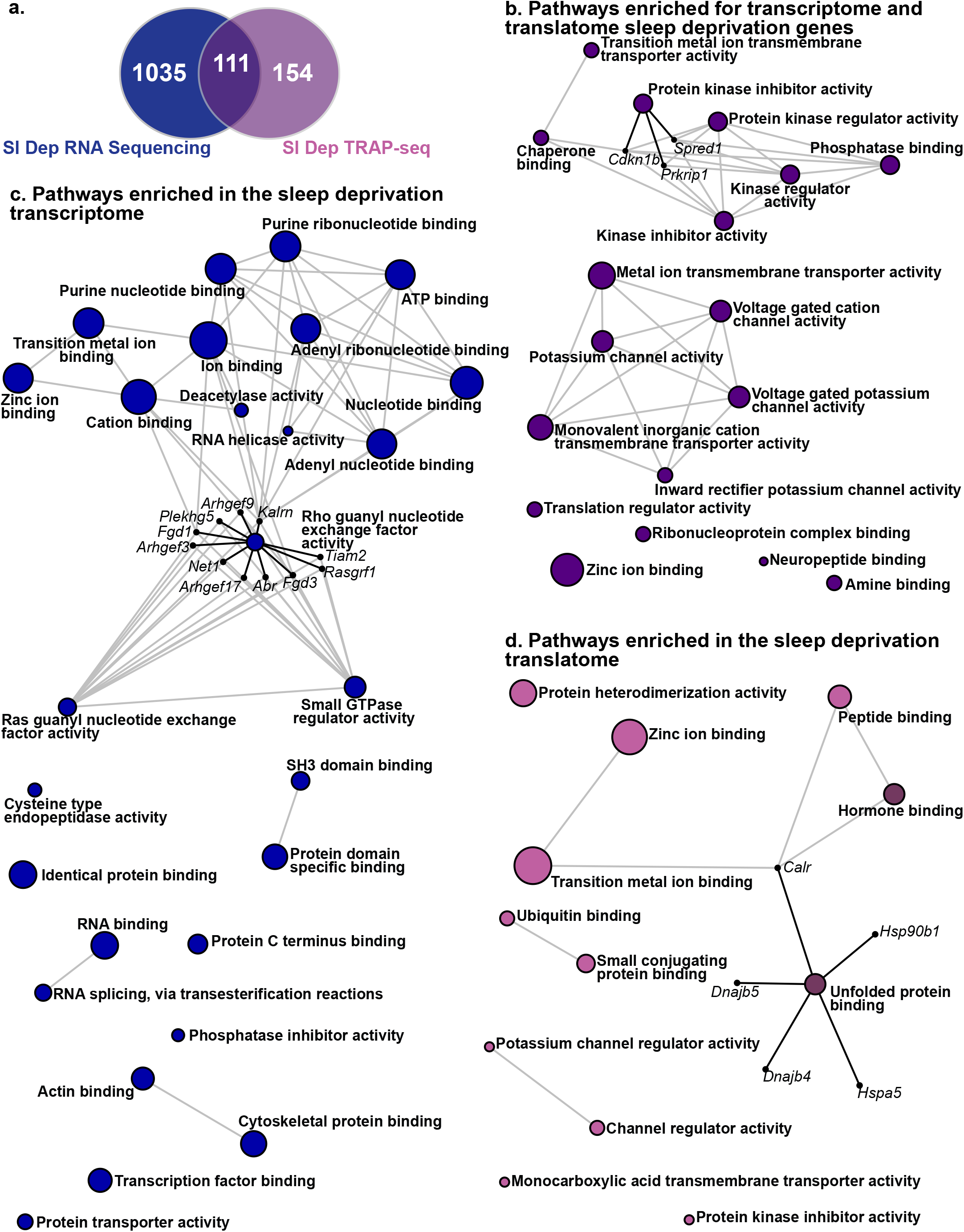
Acute sleep deprivation causes distinct transcriptome and translatome patterns. **(a)** We compared the top RNA-seq genes (1,146 genes) to our previously generated TRAP-Seq sleep deprivation results (265 genes) and identified 1,035 genes only identified as differentially expressed using RNA-seq (transcriptome), 154 genes only identified as differentially expressed using TRAP-Seq (translatome), and 111 genes found in both. **(b)** Using Network Analyst software and the Gene Ontology (GO): Molecular Function (MF) classification, pathways enriched for genes found using both RNA-seq and TRAP-Seq. The most significant pathway was Protein kinase inhibitor activity (P-value = 2.01 × 10^−3^, Adjusted P-value = 0.366) and this network has been expanded to show the genes that are transcribed and translated after sleep deprivation. **(c)** Using Network Analyst software and the GO:MF classification, pathways enriched for genes found using only RNA sequencing. The most significant pathway was Rho guanyl nucleotide exchange factor activity (P-value = 3.07 × 10^−4^, Adjusted P-value = 0.119) and this network has been expanded to show the genes that are involved and transcribed after sleep deprivation. **(d)** Using Network Analyst software and the GO:MF classification, pathways enriched for genes found using only TRAP-Seq. The most significant pathway was Unfolded protein binding (Adjusted P-value = 0.021) and this network has been expanded to show the genes that are involved and translated after sleep deprivation. Networks that survive correction for multiple testing (Adjusted P-value < 0.05) are emphasized with dark pink. The size of each node represents the number of hits from the inputted gene list.

To provide additional insight into the impact of sleep deprivation on the transcriptome and translatome, we performed pathway analyses using GO:MF for genes commonly regulated at both levels, genes regulated solely in the transcriptome, and genes that were uniquely regulated in the translatome. Pathway analysis of the genes regulated similarly after sleep deprivation in the transcriptome and the translatome revealed two major groups of gene enrichment (Figure 6b and Additional File 2: Table S9). The first group of enriched pathways included protein kinase and protein phosphatase pathways suggesting that sleep deprivation strongly affects cell signaling pathways. Secondly, there was enrichment of potassium channel activity pathways suggesting that sleep deprivation affects neural activity through gene regulation. Analysis of genes regulated solely in the transcriptome highlighted pathways involved in transcription factor binding, histone deacetylase activity, nucleotide binding, nucleotide exchange factor activity and small GTPase regulator activity (Figure 6c and Additional File 2: Table S10). In contrast, the translatome enrichment networks included the unfolded protein binding pathway, protein binding, peptide binding, protein dimerization, and ubiquitin binding (Figure 6d and Additional File 2: Table S11). This comparative analysis highlights the multiple levels of gene regulation impacted by sleep deprivation with distinct consequences.

## Discussion

Numerous behavioral and molecular studies have demonstrated the requirement of sleep for memory and neural plasticity (reviewed in [53, 54]). Previous research has shown that the hippocampus is highly susceptible to the effects of acute sleep deprivation inhibiting long-term memory with changes apparent in neuronal connectivity and morphology [14, 20, 22, 55, 56]. Although studies using mice and rats have shown that acute sleep deprivation affects gene expression in the hippocampus and the forebrain [25, 48, 52, 57-62], much of the previous research has focused on specific gene sets or used microarray analysis, rendering an incomplete picture of the effects of sleep deprivation on gene expression. Thus, we conducted an unbiased investigation of the effects of sleep deprivation on hippocampal gene expression using RNA-Seq, As predicted, the RNA-Seq experiments provided a more in-depth investigation into the effects of sleep deprivation on the transcriptome with more genes analyzed than previous microarray experiments. We found that five hours of sleep deprivation upregulated or downregulated gene expression dependent upon the biological functions and cellular components associated with the genes. The RNA-Seq results were validated through independent sleep deprivation and recovery experiments followed by RT-qPCR for genes of interest.

Analysis of genes upregulated by sleep deprivation revealed associations with nuclear functions including genes involved in RNA binding, processing, and splicing potentially increasing RNA splicing misregulation, nonsense mediated decay and RNA degradation. For example, the *Upf2* gene, a mediator of nonsense mediated decay, was upregulated by sleep deprivation consistent with the hypothesis that sleep deprivation could result in changes in RNA splicing that lead to increased RNA degradation. Misregulation of RNA splicing affects neural plasticity and function (reviewed in [48]). Dysregulation of RNA binding proteins and splicing has been associated with aberrant neural function and neurodegenerative diseases including Alzheimer’s disease (reviewed in [63, 64]). Thus, acute sleep deprivation has the potential to induce widespread changes in neuronal and synaptic plasticity through changes in RNA processing.

In the present study, we found that significant downregulation of genes by sleep deprivation was associated with cell adhesion and synaptic protein functions including *Nlgn1, Nlgn3, Ncam1, Nectin3* and *Nectin4*. Cell adhesion molecules, such as the post-synaptic adhesion protein Neuroligin-1 has been previously associated with sleep regulation ([62] and reviewed in [65]). Sleep deprivation downregulated metalloproteases such as *Adam23*, involved in cell-cell interactions. Although multiple cellular components were significantly enriched for genes downregulated by sleep deprivation, the top three cellular locations were the dendrite, postsynaptic membrane, and the synapse. Postsynaptic density scaffolding proteins, such as members of the Disc large associated protein family *Dlgap1* and *Dlgap3*, were significantly downregulated by sleep deprivation. Thus, the probable outcomes of the downregulation of genes by sleep deprivation are weakened synaptic plasticity and cell-cell interactions. Our results are consistent with previously observed decreases in hippocampal plasticity seen following brief periods of sleep deprivation [66-69]. Sleep deprivation appears to have some of the largest cellular impacts at the synapse as recent studies on whole forebrain tissue found that acute sleep deprivation reduced the rhythms in protein phosphorylation of synaptic proteins [70, 71]. Although differences in the effect of sleep deprivation on transcription and translation are apparent between brain regions ([48, 51] and reviewed in [5]), acute sleep deprivation also affects synaptic proteins in the cortex [72].

Three hours of recovery sleep following acute sleep deprivation normalized gene expression for most genes we investigated, similar to what has been observed for many genes in the hippocampus and the cortex [25, 52]. However, we did find that the transcription factor *Nr4a1* remained upregulated after recovery sleep, albeit to a smaller extent. In addition, the RNA splicing factor *Srsf7* reversed direction showing a significant decrease in expression after recovery sleep. These results indicate that recovery from sleep deprivation is gene specific, rather than a universal return of gene expression to baseline levels. Previously, expression of the transcription factor Elk1 in the hippocampus was shown to remain high after 2.5 hours of recover sleep [25]. Studies in the cortex found that some genes normalize expression levels quickly with recovery sleep, while other genes require up to 6 hours of recovery sleep to return to baseline levels [52]. Of note, genes that responded less quickly to recovery sleep included genes involved in RNA splicing and RNA binding proteins [52]. Thus, in addition to the more immediate effects of acute sleep deprivation on gene expression and the subsequent impact on synaptic plasticity and cellular signaling, acute sleep deprivation may also exert longer lasting effects on gene regulation through the continued dysregulation of transcription factors and genes related to RNA processing. Further cell-specific research needs to be completed to fully investigate the persistent effects of acute sleep deprivation on RNA splicing and processing.

The canonical view of gene regulation and the central dogma of molecular biology suggest that RNA and protein abundance are highly correlated. However, as understanding of RNA processing increased, it has become apparent that gene regulation occurs at multiple levels (reviewed in [73]). Acute sleep deprivation, in particular, appears to distinctly impact transcription and translation as we found when we compared the results of the current RNA-Seq data set with the translatome of excitatory neurons in the hippocampus after sleep deprivation. Although differences arose between these data sets due to cell type differences, genes impacted similarly by sleep deprivation in the transcriptome and translatome encompassed genes involved in kinase and phosphatase signaling pathways, as well as cation and potassium channels. Changes in the expression of membrane channels and cellular signaling pathways have the potential to rapidly impact synaptic strength and plasticity following sleep deprivation. A large number of genes, more than 1,000, were upregulated in the transcriptome, but not in the translatome. Genes regulated only at the level of the transcriptome included genes involved in RNA processing, nucleotide binding and small GTPase signaling. Potentially genes upregulated in the transcriptome, but not the translatome, may reflect transcripts with alternative splicing that are not translated efficiently or that undergo degradation. Alternatively, genes upregulated in the transcriptome by sleep deprivation may also be sequestered in dynamic RNA granules that may then be released when conditions normalize (reviewed in [74-76]). Notably, there are also genes downregulated in the transcriptome but not in the translatome. Potentially, the translatome reflects isoform specific transcript association with the ribosome, while our RNA-Seq data set reflects total RNA abundance. A comparatively small number of genes, approximately 150 genes, showed separate regulation by sleep deprivation in the translatome of excitatory neurons but not in the overall transcriptome including genes associated with the unfolded protein response and ubiquitination suggesting that additional regulation of protein degradation may occur with sleep deprivation. The analyses presented provide further insight into the nuanced effects of sleep deprivation on gene regulation at multiple levels.

The results presented here provide an unbiased in-depth perspective of the effects of acute sleep deprivation on gene expression in the hippocampus. Notably, our results clearly demonstrate that sleep deprivation differentially upregulates or downregulates genes dependent upon biological function, instead of a more general mechanism resulting in global changes in gene expression. Moreover, our analyses provided new insight into the effects of sleep deprivation revealing the strong association of genes upregulated by sleep deprivation with nuclear functions. In contrast, genes downregulated by sleep deprivation were associated with multiple cellular components, in particular, dendrites and synapses. These distinctions highlight the need for future research investigating the effects of sleep deprivation on the hippocampus taking advantage of technological advances in single cell and spatial transcriptomics. Although the results presented here establish a strong foundation for comparison with data from other brain regions to more precisely understand brain region specific impacts of sleep deprivation, further research is needed to understand the persistent effects of acute sleep deprivation on long-term memory, as well as to identify the effects of chronic sleep restriction on the hippocampus.

## Declarations

### Ethics approval and consent to participate

Not applicable

### Consent for publication

Not applicable

### Availability of data and materials

The datasets generated and/or analyzed during the current study are available in the NCBI’s Gene Expression Omnibus repository, GEO Series accession GSE166831, https://www.ncbi.nlm.nih.gov/geo/query/acc.cgi?acc=GSE166831; GEO accession GSE156925, https://www.ncbi.nlm.nih.gov/geo/query/acc.cgi?acc=GSE156925 and GEO accession GSE33302, https://www.ncbi.nlm.nih.gov/geo/query/acc.cgi?acc=GSE33302.

### Authors’ contributions

LCL, MEG and TA conceived the study. LCL and MEG planned the experiments, performed sleep deprivation, RT-qPCR and data analysis. SC extracted the RNA for library preparation. EB performed the RNA sequencing bioinformatic analyses with review by JJM. All authors read and approved the final manuscript.

## Acknowledgements

Not applicable

## Additional Information

### Additional File 1

**Figure S1. Normalization of RNA sequencing data**. Distributional differences in GC content and variability in sequencing depth are sources of technical variability in RNA Sequencing data. GC content distributions **(a)** before normalization, **(b)** after full quantile GC content normalization, and **(c)** upper quartile sequencing depth normalization. Relative log expression (RLE) plots **(d)** before normalization, **(e)** after full quantile GC content normalization, and **(f)** upper quartile sequencing depth normalization. Principal component analysis (PCA) plots **(g)** before normalization, **(h)** after full quantile GC content normalization, and **(i)** upper quartile sequencing depth normalization.

**Figure S2. Validation of negative controls**. From an independent cohort of mice (n=6 in each group), RT-qPCR was used to validate the findings of negative control genes that showed no differential expression in the RNA sequencing results. **(a)** *Lama5* (P-value = 0.9393), **(b)** *Fzd5* (P-value = 0.1573), and **(c)** *Trpm3* (P-value = 0.5076). Data are presented as mean ± SEM and normalized against two housekeeping genes (*Tubulin* and *Hprt*). Comparisons are evaluated using an unpaired t-test and n.s. denotes non-significant differences (P > 0.05).

### Additional File 2

**Table S1: RT-qPCR primers used**. Validation of RNA-seq results was performed using RT-qPCR, SYBR technology, and custom designed primers.

**Table S2: Genes significantly upregulated by sleep deprivation**. We compared non-sleep deprived and sleep deprived mice and identified 507 genes with a false discovery rate (FDR) < 0.1 that were upregulated after sleep deprivation. Gene types include antisense, bidirectional promoter long non-coding RNA (lncRNA), long intergenic non-coding RNA (lincRNA), processed transcript, protein coding, Small Cajal body-specific RNA (scaRNA), to be experimentally confirmed (TEC), transcribed processed pseudogene, transcribed unprocessed pseudogene, unitary pseudogene, and unprocessed pseudogene. LogFC denotes Log Fold Change, LogCPM denoted Log Counts Per Million, and F denotes the F-statistic from edgeR’s quasi-likelihood pipeline.

**Table S3: Enrichment networks in upregulated genes using PANTHER:BP**. We used Network Analyst software and the PANTHER:Biological Processes (BP) classification to perform overrepresentation analysis (ORA) and identified 16 pathways enriched for upregulated genes. Hits represent the number of upregulated genes found in each network.

**Table S4: Cellular components enriched in upregulated genes using Panther:CC**. We used Network Analyst software and the PANTHER: Cellular Components (CC) classification to perform overrepresentation analysis (ORA) and identified 10 networks enriched for upregulated genes. Hits represent the number of upregulated genes found in each network.

**Table S5: Genes significantly downregulated by sleep deprivation**. We compared sleep deprived and undisturbed mice and identified 639 genes with an FDR < 0.1 that were downregulated after sleep deprivation. Gene types include antisense, bidirectional promoter lncRNA, lincRNA, microRNA, processed pseudogene, processed transcript, protein coding, sense intronic, TEC, transcribed processed pseudogene, and transcribed unprocessed pseudogene. LogFC denotes Log Fold Change, LogCPM denoted Log Counts Per Million, and F denotes the F-statistic from edgeR’s quasi-likelihood pipeline.

**Table S6: Enrichment networks in downregulated genes using PANTHER:BP**. We used Network Analyst software and the PANTHER:BP classification to perform ORA and identified 19 pathways enriched for downregulated genes. Hits represent the number of downregulated genes found in each network.

**Table S7: Cellular components enriched in downregulated genes using PANTHER:CC**. We used Network Analyst software and the PANTHER:CC classification to perform ORA and identified 19 networks enriched for upregulated genes. Hits represent the number of downregulated genes found in each network.

**Table S8: Comparison of differentially expressed genes after sleep deprivation between whole hippocampal transcriptome and TRAP-Seq from excitatory neurons**. To identify the sleep deprivation transcriptome and translatome we compared RNA-seq and TRAP-Seq datasets. The first column shows 1,035 differentially expressed genes only found using RNA sequencing, the second column shows 111 differentially expressed genes found in both RNA sequencing and TRAP-Seq, and the third column shows 154 differentially expressed genes only found using TRAP-Seq. Lists are sorted alphabetically.

**Table S9: Molecular function networks enriched in genes significantly affected by sleep deprivation in RNA-seq and TRAP-Seq data sets using GO:MF**. We used Network Analyst software and the Gene Ontology (GO): Molecular Function (MF) classification to perform ORA and identified 18 networks enriched for genes differentially expressed using RNA-seq and TRAP-Seq. Hits represent the number of genes found in each network.

**Table S10: Molecular function networks for genes differentially regulated by sleep deprivation only in whole hippocampal transcriptome using GO:MF**. We used Network Analyst software and the Gene Ontology (GO): Molecular Function (MF) classification to perform ORA and identified 27 networks enriched for genes only differentially expressed using RNA sequencing. Hits represent the number of genes found in each network.

**Table S11: Molecular function networks for genes differentially regulated by sleep deprivation only in the TRAP-Seq data set using GO:MF**. We used Network Analyst software and the GO:MF classification to perform ORA and identified 12 networks enriched for genes differentially expressed using only TRAP-Seq. Hits represent the number of genes found in each network.

