## Supplementary figures and images for "Altered hippocampal transcriptome dynamics following sleep deprivation"

### Supplemental Figures

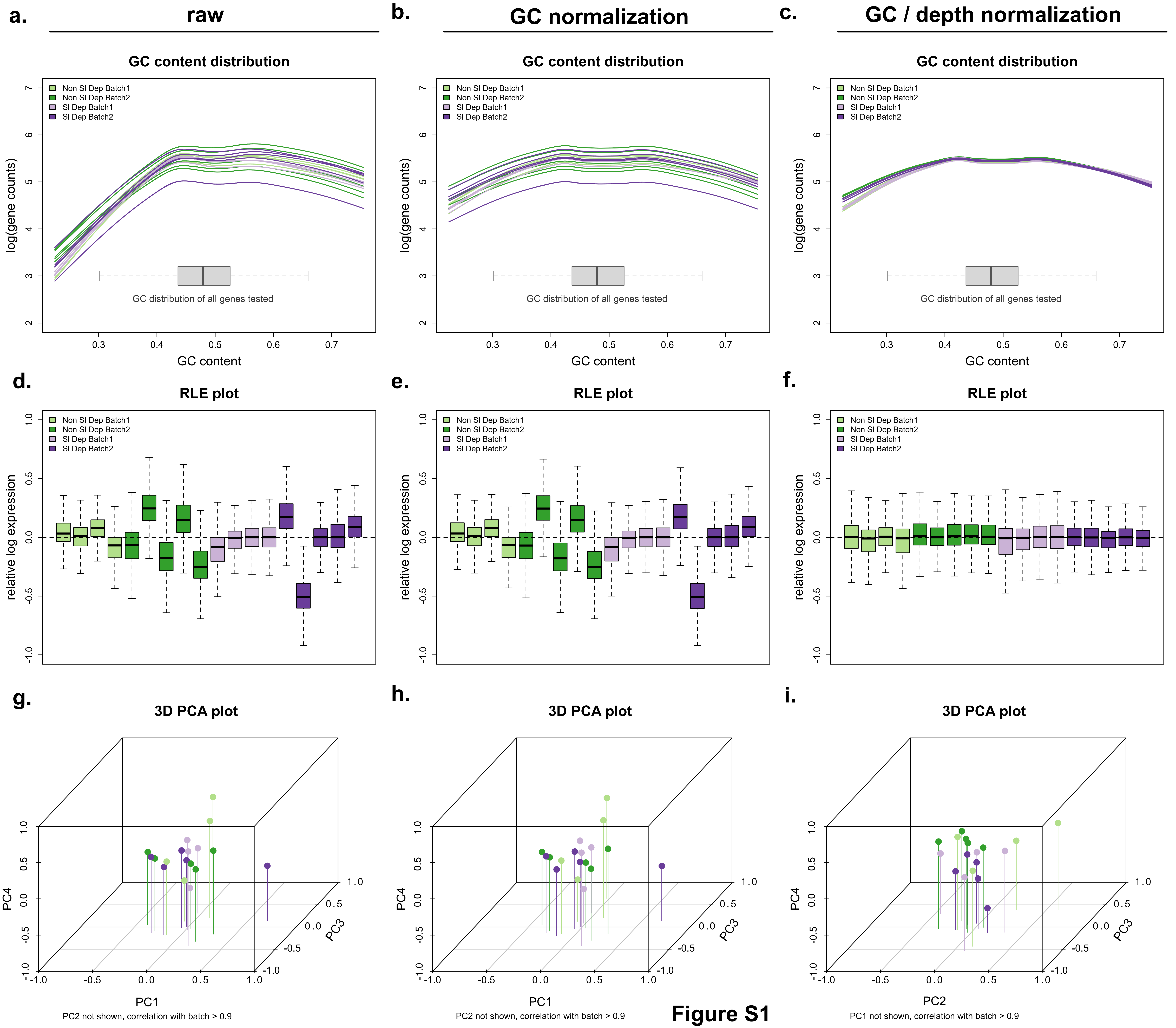

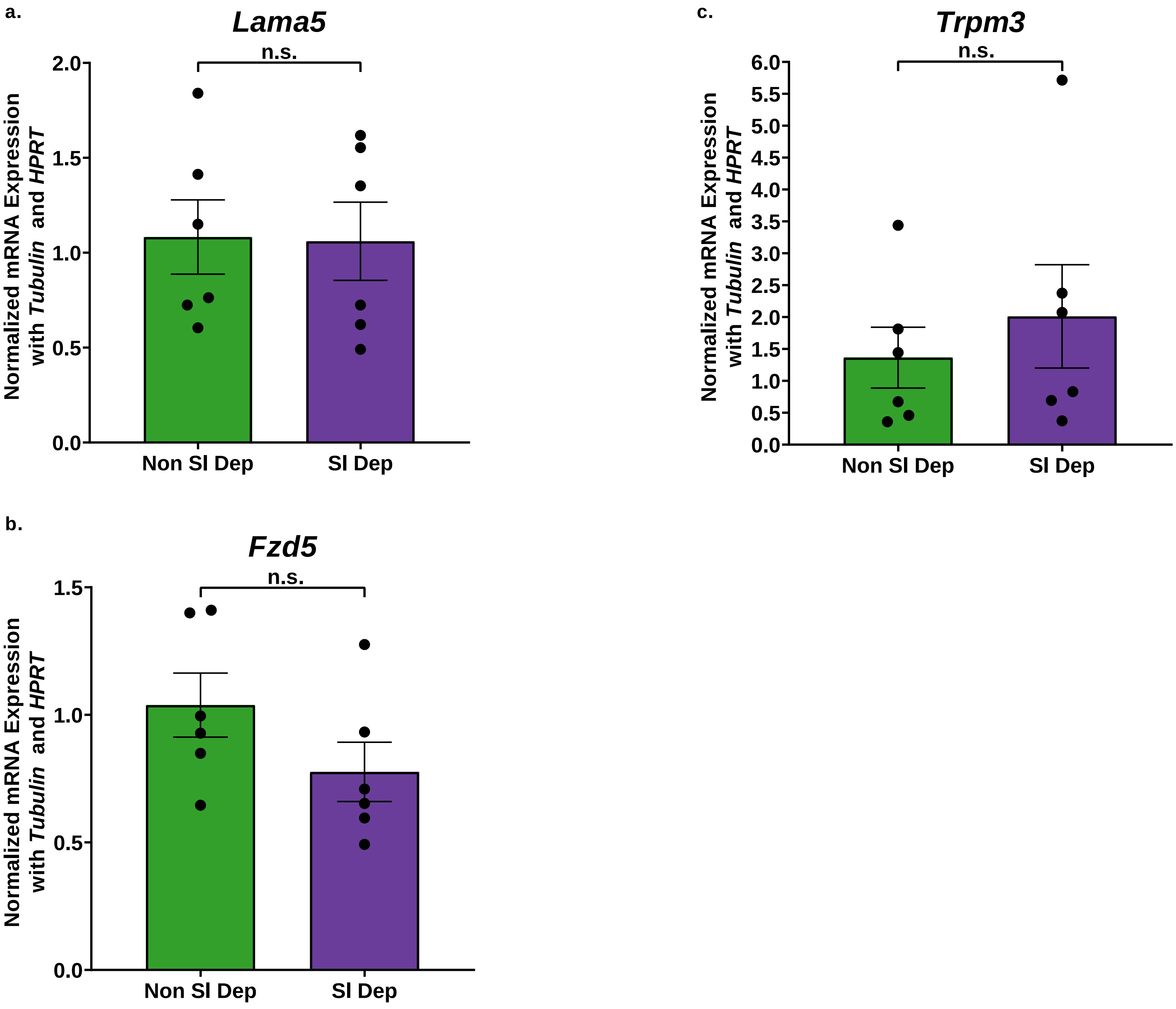
